# Mapping of Stripe Rust and Leaf Rust Resistance Genes in the Hard Red Winter Wheat Population Green Hammer × Lonerider

**DOI:** 10.64898/2026.05.13.724876

**Authors:** Rajat Sharma, Meinan Wang, Xianming Chen, Brett F. Carver, Mary Guttieri, Paul St. Amand, Amy Bernardo, Guihua Bai, Shuyu Liu, Anju Maan Ara, Meriem Aoun

## Abstract

Stripe rust and leaf rust, caused by *Puccinia striiformis* f. sp. *tritici* and *P. triticina*, respectively, are the most destructive wheat diseases in the southern Great Plains. ‘Green Hammer’ is a hard red winter wheat (HRWW) cultivar released by Oklahoma State University in 2018 and has demonstrated a stable adult plant resistance to stripe rust and race-specific seedling resistance to leaf rust. To identify and map rust resistance loci, 109 doubled haploid (DH) lines derived from the cross between Green Hammer and another HRWW cultivar, ‘Lonerider’, were developed. Lonerider showed adult plant resistance to stripe rust but was susceptible to multiple *P. triticina* races. The DH lines were evaluated for stripe rust at the adult plant stage in greenhouse and field environments across Oklahoma, Kansas, and Washington, and for leaf rust at the seedling stage against seven U.S. *P. triticina* races and at the adult plant stage in Oklahoma and Texas. Genotyping-by-sequencing generated 6,078 polymorphic single-nucleotide polymorphisms used for genetic mapping. Quantitative trait loci (QTL) analysis identified 14 stripe rust and 8 leaf rust resistance QTL. For stripe rust, a major QTL in Green Hammer, *QYr.osughln-2AS*, was identified in the proximity of the 2N^v^S translocation. Three other major stripe rust resistance QTL were identified in Lonerider on chromosomes 2AL (two QTL) and 2BS (one QTL). For leaf rust, *QLr.osughln-1DS* and *QLr.osughln-2DS.1* were the two major QTL identified in Green Hammer and most likely correspond to the all-stage resistance genes *Lr21* and *Lr39*, respectively. In this study, we identified previously characterized genes as well as unknown genes that can be utilized in wheat breeding programs to enhance resistance to leaf rust and stripe rust.

## Introduction

Stripe rust and leaf rust, caused by the biotrophic fungi *Puccinia striiformis* f. sp. *tritici* (*Pst*) and *P. triticina* (*Pt*), respectively, are among the most devastating diseases of wheat worldwide. Globally, leaf rust is considered the most common wheat rust, and stripe rust poses a threat to approximately 88% of wheat-growing acreage worldwide (Beddow et al. 2015; Bolton et al. 2008). Leaf rust and stripe rust can cause yield losses of up to 40% and 100%, respectively, under favorable conditions (Chen 2005; Li et al. 2014). Historically, *Pst* was adapted to cool, moist conditions typical of the Pacific Northwest (PNW) of the United States (US). However, the emergence of new races after the 2000s has resulted in populations that are better adapted and more aggressive under warmer temperature conditions of the US Great Plains (Chen et al. 2010; Markell and Milus 2008; Milus et al. 2009). *Pt* has a temperature optimal window of 15–25°C and exhibits high virulence diversity in the US, with approximately 30–60 races identified annually (Kolmer and Fajolu 2022). Although wheat rusts can be managed using foliar fungicides, this approach increases production costs and raises concerns about environmental negative effects. Therefore, cultivation of resistant cultivars is a preferred management strategy because it is cost-effective and environmentally friendly (Chen 2005).

Rust resistance genes, including stripe rust resistance genes (*Yr*) and leaf rust resistance genes (*Lr*), are classified into two types based on their effectiveness and the growth stage at which they confer resistance, namely all-stage resistance (ASR) or seedling resistance and adult plant resistance (APR). To date, 87 *Yr* genes and *85 Lr* genes have been permanently designated (McIntosh et al. 2020; Sharma et al. 2024). Of the 87 *Yr* genes, 60 confer ASR, whereas the remaining 27 confer APR (Sharma et al. 2026). Similarly, among the 85 *Lr* genes, most confer ASR, with only 8 genes conferring APR, namely *Lr34*/*Yr18*, *Lr46*/*Yr29*, *Lr67*/*Yr46*, *Lr68*, *Lr74*, *Lr75*, *Lr77*, and *Lr78* (Lakkakula et al. 2025).

Among officially designated *Yr* and *Lr* genes, 7 *Yr* ASR genes and 11 *Lr* ASR genes have been cloned, including *Yr5*/*Yr7*/*YrSP* (Marchal et al. 2018), *Yr10* (*YrNAM*) (Liu et al. 2014; Ni et al. 2023), *Yr15* (Klymiuk et al. 2018), *Yr27* (Athiyannan et al. 2022), *Yr84* (Klymiuk et al. 2025), *Yr87*/*Lr85* (Sharma et al. 2024), *Lr1* (Cloutier et al. 2007), *Lr9* (Wang et al. 2023), *Lr10* (Feuillet et al. 2003), *Lr13* (Hewitt et al. 2021; Yan et al. 2021), *Lr14a* (Kolodziej et al. 2021), *Lr21* (Huang et al. 2003), *Lr22a* (Thind et al. 2017), *Lr30* (Yang et al. 2025), *Lr42* (Lin et al. 2022), and *Lr47* (Li et al. 2023a). Only three APR *Lr*/*Yr* genes, including *Lr34*/*Yr18* (Krattinger et al., 2009), *Yr36* (Fu et al., 2009), and *Lr67*/*Yr46* (Moore et al., 2015), have been cloned.

All-stage resistance is typically race-specific and conferred by major resistance genes that are expressed across all plant growth stages. They often encode nucleotide-binding leucine-rich repeat (NLR) proteins that act as receptors for specific pathogen avirulence effector and trigger a defense response. ASR commonly follows the gene-for-gene concept and is short-lived because it imposes strong selection pressure on the pathogen, promoting the emergence of new virulent races. In contrast, APR genes are generally more durable, often race non-specific, and confer a slow-rusting resistance at the adult plant stage. The cloned *Yr* and *Lr* APR genes *Lr34*/*Yr18*, *Yr36*, and *Lr67*/*Yr46* encode a putative ATP-binding cassette transporter (Krattinger et al. 2009), a protein kinase (WKS1) (Fu et al. 2009), and a hexose transporter (Moore et al. 2015), respectively. However, a single APR gene typically confers partial resistance and may be insufficient under severe epidemics unless multiple APR genes are pyramided (Chen 2014; Risk et al. 2012; Singh et al. 2015; Sørensen et al. 2014). Therefore, pyramiding multiple APR genes or in combination with effective ASR genes is needed to achieve high and durable rust resistance (Sharma et al. 2025). Characterizing rust resistance genes in modern wheat cultivars is important for their efficient deployment in breeding programs, particularly because these genes often carry minimal linkage drag.

‘Green Hammer’ (pedigree: OK Bullet/TX00D1390//Shocker; experimental line: OK13209) is a hard red winter wheat (HRWW) cultivar released in 2018 by the Oklahoma State University (OSU) wheat breeding program. In recent years, Green Hammer has been the second most planted cultivar in Oklahoma, likely due to its favorable combination of disease resistance, good baking and milling quality, and high yield. Since its early development in 2013, Green Hammer has shown high to moderate but stable APR to stripe rust in the Great Plains. For leaf rust, Green Hammer confers race-specific seedling resistance and stable and high APR in wheat fields in the southern Great Plains. However, the genetic basis of stripe rust and leaf rust resistance in Green Hammer is not fully understood. Therefore, the objective of this study was to dissect and characterize stripe rust and leaf rust resistance in Green Hammer using a doubled haploid (DH) population derived from the bi-parental cross Green Hammer × Lonerider.

## Materials and Methods

### Plant materials

A total of 109 DH lines developed from cross Green Hammer × Lonerider were used to map quantitative trait loci (QTL) for stripe rust and leaf rust resistance. Green Hammer was evaluated in the Southern Regional Performance Nursery (SRPN) in 2017 and was found to be susceptible at the seedling stage to major U.S. *Pst* races, including PSTv-37 (https://www.ars.usda.gov/plains-area/lincoln-ne/wheat-sorghum-and-forage-research/docs/hard-winter-wheat-regional-nursery-program/research/). PSTv-37 is the predominant race and is virulent to *Yr6*, *Yr7*, *Yr8*, *Yr9*, *Yr17*, *Yr27*, *Yr43*, *Yr44*, *Yr85* (= *YrTr1*), and *YrExp2* (Wan and Chen 2014; Wang et al. 2022). However, Green Hammer has shown consistent APR to stripe rust and leaf rust and also carries race-specific leaf rust ASR. ‘Lonerider’ (pedigree: Billings/OK08328; experimental line: OK12DP22002-042) is another HRWW cultivar released by OSU in 2017. Lonerider was evaluated in SRPN in 2016 and 2017 and was susceptible at the seedling stage to major *Pst* races and to most US *Pt* races. At the adult plant stage, Lonerider ranged from moderately resistant to moderately susceptible to stripe rust across most field environments in the US. ‘Jagalene’, ‘OK Bullet’, ‘Pete’, ‘PS 279’, and ‘Thatcher’ were included as susceptible checks in phenotyping experiments.

### Stripe rust evaluation at the adult plant stage

The parents (Green Hammer and Lonerider), 109 DH lines, and susceptible checks (Pete and Jagalene) were evaluated at the flag-leaf stage under controlled greenhouse conditions using race PSTv-37, following protocols described by Sharma et al. (2026). Briefly, six plants per DH line were evaluated in three replications, with two plants per replication. Plants were vernalized at 4°C under a 12 h photoperiod for 6–8 weeks. After vernalization, plants were grown at 22°C/18°C (day/night) with a 16 h photoperiod. At the flag-leaf stage, plants were inoculated with a spore suspension of PSTv-37 urediniospores in Soltrol 170 mineral oil (Chevron Phillips Chemical Co., Woodlands, TX( at a concentration of 10 mg mL^−1^. After inoculation, plants were kept in a dark chamber at 10°C and 100% relative humidity for 24 h and then transferred to a growth chamber maintained at a diurnal cycle of 4–20°C with 16 h photoperiod. At 18–21 days post inoculation, infection type (IT) was recorded for each plant using a 0–9 scale, where 0–3 were considered highly resistant, 4–6 moderately resistant, and 7–9 susceptible (Line and Qayoum 1992; Wan et al. 2017). The mean IT of six plants across three replications for each line was used for further analysis.

The DH lines, along with the parents and susceptible checks, were also evaluated for stripe rust responses in field environments in the US Great Plains at Chickasha, Oklahoma in 2024, at Rossville, Kansas in 2024 and 2025, and in the PNW at Pullman and Mt. Vernon, Washington in 2024 and 2025. These locations differ in *Pst* race composition and weather conditions. Standard management practices were adopted across all nurseries. In Chickasha and Rossville, nurseries were artificially inoculated with local isolates with PSTv-37 as the predominant race. In Pullman and Mt. Vernon, nurseries were naturally infected, except Pullman 2025, with PSTv-37 being the most predominant race, but multiple other races were also present based on stripe rust annual surveys. The 2025 Pullman field was artificially inoculated at the early jointing stage (Feekes stage 5) with urediniospores (mostly race PSTv-37) collected from the same site in 2024. Across all field nurseries in both years, IT was recorded using a 0–9 scale, and disease severity (DS) was recorded as the percentage of infected leaf area (Peterson et al. 1948). In Chickasha in 2024, the DH lines were planted in 1.2 m rows. The parents and susceptible checks (Pete and Jagalene) were planted after every 50 DH lines. Pete was also planted as spreader and border rows. In Rossville in 2024 and 2025, the parents and susceptible checks were planted after every 80 rows. In Chickasha and Rossville, IT and DS were recorded on the flag leaves at Feekes growth stage 10.5 (Large 1954). In Pullman and Mt. Vernon, Jagalene and PS 279 were used as susceptible checks along with the parents with PS 279 was planted after every 20 rows and as border rows. In Pullman, IT and DS were recorded at Feekes stage 10.5 in 2024 and at Feekes stage 11 in 2025. In Mt. Vernon, disease ratings were recorded twice each year. The first rating was taken at jointing, which corresponded to Feekes stage 4 in 2024 and Feekes stage 5 in 2025. The second rating was taken after heading, which corresponded to Feekes stage 10.54 in 2024 and Feekes stage 11 in 2025. Best linear unbiased estimates (BLUEs) for IT and DS across all environments were estimated using a mixed linear model implemented in the R package “lme4” (Bates et al. 2015; Vazquez et al. 2010). IT and DS ratings from field nurseries, greenhouse, and BLUEs were treated as distinct traits. Pearson correlation coefficients were calculated for each stripe rust response trait using the R package “metan” (Olivoto and Lúcio 2020).

### Leaf rust evaluation at the seeding and the adult plant stages

The parents, Green Hammer and Lonerider, were evaluated for leaf rust responses at the seedling stage against 12 *Pt* races, including KFBJG, MCTNB, MGPSB, MHDSB, MJBJG, MNPSD, MPPSD, TBBGS, TCGJG, TCRDG, TFTSB, and TNBJS, to identify races for which the parents showed contrasting reactions. The virulence and avirulence profiles of these *Pt* races were determined at the seedling stage using a differential set of 20 *Lr* near-isogenic lines of Thatcher wheat (Supplementary Table S1) as described by Long and Kolmer (1989). Among the 12 tested races, 7 were avirulent to Green Hammer and virulent to Lonerider, including KFBJG, MCTNB, MGPSB, MHDSB, MJBJG, MNPSD, and TCRDG. These races were used to evaluate the DH population at the seedling stage using the protocol described by Lakkakula et al. (2025). Briefly, 5–7 seeds per DH line were planted in a 72-cell tray. The experiment was conducted with two replications, and Thatcher and the parents of the DH population were planted twice in each tray. Plants were grown in a rust-free greenhouse under a diurnal temperature cycle of 18–22°C with a 16 h photoperiod. A spore suspension of *Pt* urediniospores in Soltrol 170 mineral oil at a concentration of 10 mg mL^−1^ was used to inoculate seedlings when the first leaves had fully expanded. After inoculation, seedlings were kept in a dark chamber at 18°C and 100% relative humidity for 16–18 h. Inoculated seedlings were then returned to the greenhouse and evaluated at 10–12 days post-inoculation using the 0–4 IT scale (Stakman et al. 1962), where IT < 3 indicated resistance and IT ≥ 3 indicated susceptibility. Variation in uredinia size relative to normal was indicated using “+” and “−”. Infection types scored on the 0–4 scale were converted to a linearized 0–9 scale as described by Zhang et al. (2014), where 0–3 were considered highly resistant, 4–6 moderately resistant, and 7–9 susceptible.

The DH population was also evaluated at the adult plant stage in field environments at Chickasha and Lahoma in Oklahoma, and Castroville in Texas, in 2025. The DH lines were planted in two replications. Susceptible checks (Jagalene and OK Bullet) and the parents were planted after every 30 rows. In Chickasha and Lahoma, OK Bullet was also planted as spreader and border rows and nurseries were artificially inoculated with local *Pt* isolates collected from recent years in the field to ensure uniform and high disease pressure. Infection response and DS ratings were recorded at Feekes stage 11 (ripening). Infection response categories were based on visible symptoms, where immune indicated no visible symptoms, R (resistant) indicated necrosis or chlorosis without uredinia, MR (moderately resistant) indicated necrosis or chlorosis with small uredinia, MS (moderately susceptible) indicated medium-sized uredinia without necrosis but with possible chlorosis, and S (susceptible) indicated large uredinia without necrosis or chlorosis (McIntosh et al. 1995; Roelfs et al. 1992). Infection response and DS were combined to calculate the coefficient of infection (COI), which was computed as the product of DS and a constant value assigned to infection response, where immune = 0.0, R = 0.2, RMR = 0.3, MR = 0.4, MRMS = 0.5, MSMR = 0.6, MS = 0.8, MSS = 0.9, and S = 1.0 (Aoun et al. 2016; Yu et al. 2011). In Castroville, the leaf rust nursery was naturally infected with local Texas *Pt* isolates and leaf rust responses were evaluated at the adult plant stage using a disease response (DR) scale of 0–9 as described by Lakkakula et al. (2025). For all environments, mean leaf rust responses across replications for each line were used for further analysis.

### Genotyping, linkage map construction, and QTL analysis

Genomic DNA was extracted from young seedlings at the two-leaf stage, and genotyping was conducted using genotyping-by-sequencing (GBS) at the USDA-ARS Genotyping Lab in Manhattan, KS, as described by Sharma et al. (2026). Single-nucleotide polymorphism (SNP) calling and SNP alignment were performed using TASSEL v5 (Bradbury et al. 2007) and bwa samse v0.7.17 (Li and Durbin 2009), respectively. Physical positions of SNP markers were assigned using the Chinese Spring reference genome IWGSC RefSeq v2.1 (Zhu et al. 2021). GBS initially generated 57,353 SNP markers with missing data up to 80%, minor allele frequency ≥ 5%, and heterozygosity ≤ 15%. For further analysis, 12,159 GBS-derived SNP markers with missing data ≤ 40% were retained (Supplementary Table S2). Monomorphic SNP markers as well as those showing missing data or heterozygous SNP calls in either parent, were removed. The remaining 6,078 polymorphic markers were used for QTL analysis (Supplementary Table S3).

Linkage map construction and QTL analysis were performed in QTL IciMapping v4.2 (Meng et al. 2015) as described by Sharma et al. (2026). Briefly, the 6,078 high-quality SNP markers were filtered for redundancy (same genetic position) and segregation distortion (1:1 ratio at *p* ≤ 0.01) using the BIN function. Markers were then checked for excessive crossover counts, double crossovers, and switch alleles using the R package “R/qtl” (Broman 2010). The linkage map was constructed using a minimum logarithm of odds (LOD) score of 6.0 with the MAP function in QTL IciMapping v4.2. Markers were assigned to chromosomes based on their physical positions on the Chinese Spring reference genome IWGSC RefSeq v2.1 (Zhu et al. 2021). The Kosambi mapping function was used to convert recombination frequencies into genetic distances in centiMorgans (cM) (Kosambi 1943).

Phenotypic data for stripe rust and leaf rust responses across environments, together with the linkage map, were used for QTL detection using the BIP function. Inclusive composite interval mapping for additive QTL was conducted with a walk speed of 1 cM and a stepwise regression probability of 0.001. Significant QTL were detected based on LOD thresholds determined using 1,000 permutations at *p* = 0.05. For stripe rust APR, QTL were considered major if they were detected in at least two environments and explained ≥ 10% of the phenotypic variation. For leaf rust, QTL explaining ≥ 10% of the phenotypic variation were considered major. All other QTL were considered minor.

## Results

### Stripe rust response at the adult plant stage

Green Hammer showed high resistance to moderately susceptible to stripe rust at the adult plant stage, with IT ranging from 2 (2024 Mt. Vernon 1) to 7 (2024 Pullman) and DS ranging from 5% (2024 Rossville) to 45% (2024 Pullman) (Table 1; Fig. 1). Lonerider responses at the adult plant stage ranged from highly resistant in 2024 Chickasha (IT 2, DS 7%) to moderately resistant in 2024 Pullman (IT 6, DS 40%). However, Lonerider was moderately susceptible in 2024 in Mt. Vernon at the jointing stage (IT 8, DS 60%) (Table 1; Fig. 1). Under greenhouse conditions with a low-temperature profile (4–20°C), both Green Hammer and Lonerider were highly resistant to race PSTv-37, with IT values of 3.0 and 2.5, respectively. Across environments, mean IT values of the DH lines ranged from 2.5 to 6.2, and mean DS values ranged from 6.1% to 57.2% (Supplementary Table S4). The distributions for IT and DS in the DH population showed transgressive segregation, suggesting that both parents are contributing different resistance genes (Fig. 1). In the Great Plains field environments at Chickasha and Rossville, the overall distribution of DH lines fell mostly within the highly resistant to moderately resistant categories, with only a few susceptible lines observed in 2025 Rossville (Fig. 1). Based on multi-environment BLUEs, mean IT values for Green Hammer, Lonerider, and the DH lines were 4.1, 4.0, and 4.3, and the corresponding mean DS values were 21.5%, 30.0%, and 31.5%, respectively (Table 1; Supplementary Table S4). There were significant and positive correlations between IT and DS values in most environments. However, correlations involving environment Rossville (2024), Rossville (2025), and Pullman (2025) were not significant for some environment pairs (Supplementary Fig. S1).

**Fig. 1.**
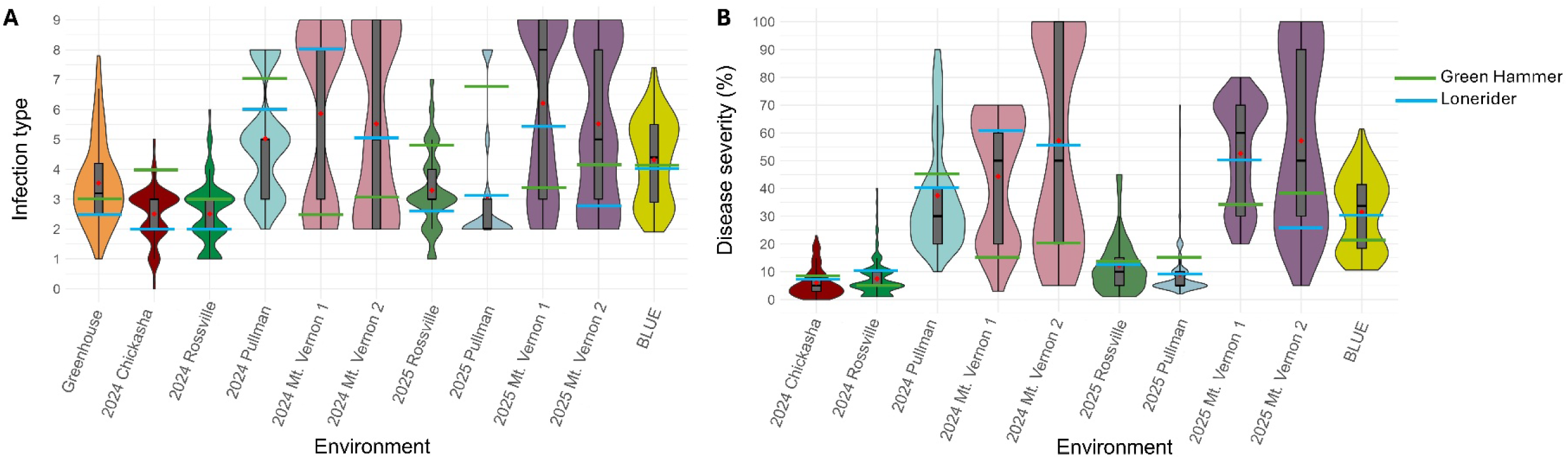
Distribution of stripe rust responses in the doubled haploid population Green Hammer × Lonerider across multiple environments. **A,** infection types and **B,** disease severity. In the boxplots, black bold horizontal lines represent the medians and red diamonds represent the means. Green and blue horizontal lines indicate the mean stripe rust responses of the parents, Green Hammer and Lonerider, respectively. BLUE = multi-environment best linear unbiased estimates.

**TABLE 1.** Stripe rust infection types and disease severity (%) for Green Hammer and Lonerider evaluated across multiple environments.

| Trait–environment<sup>a</sup> | Designation | Parents |  |
| --- | --- | --- | --- |
|  |  | Green Hammer | Lonerider |
| IT–Greenhouse (4°C–20°C) | Greenhouse-IT | 3.0 | 2.5 |
| IT–Chickasha, OK (May 4, 2024) | 2024 Chickasha-IT | 4.0 | 2.0 |
| IT–Rossville, KS (May 17, 2024) | 2024 Rossville-IT | 3.0 | 2.0 |
| IT–Pullman, WA (June 12, 2024) | 2024 Pullman-IT | 7.0 | 6.0 |
| IT–Mount Vernon (April 10, 2024) | 2024 Mt. Vernon 1-IT | 2.0 | 8.0 |
| IT–Mount Vernon (June 6, 2024) | 2024 Mt. Vernon 2-IT | 3.0 | 5.0 |
| IT–Rossville, KS (May 21, 2025) | 2025 Rossville-IT | 4.8 | 2.6 |
| IT–Pullman, WA (June 17, 2025) | 2025 Pullman-IT | 6.8 | 3.2 |
| IT–Mount Vernon (April 16, 2025) | 2025 Mt. Vernon 1-IT | 3.4 | 5.4 |
| IT–Mount Vernon (June 4, 2025) | 2025 Mt. Vernon 2-IT | 4.2 | 2.8 |
| DS–Chickasha, OK (May 4, 2024) | 2024 Chickasha-DS | 8.0 | 7.0 |
| DS–Rossville, KS (May 17, 2024) | 2024 Rossville-DS | 5.0 | 10.0 |
| DS–Pullman, WA (June 12, 2024) | 2024 Pullman-DS | 45.0 | 40.0 |
| DS–Mount Vernon (April 10, 2024) | 2024 Mt. Vernon 1-DS | 15.0 | 60.0 |
| DS–Mount Vernon (June 6, 2024) | 2024 Mt. Vernon 2-DS | 20.0 | 55.0 |
| DS–Rossville, KS (May 21, 2025) | 2025 Rossville-DS | 14.0 | 13.0 |
| DS–Pullman, WA (June 17, 2025) | 2025 Pullman-DS | 15.0 | 9.0 |
| DS–Mount Vernon (April 16, 2025) | 2025 Mt. Vernon 1-DS | 34.0 | 50.0 |
| DS–Mount Vernon (June 4, 2025) | 2025 Mt. Vernon 2-DS | 38.0 | 26.0 |
| IT–Multi-environment BLUE | BLUE-IT | 4.1 | 4.0 |
| DS–Multi-environment BLUE | BLUE-DS | 21.5 | 30.0 |

<sup>a</sup> IT = Infection type; DS = Disease severity, BLUE = Best linear unbiased estimates; Traits at Mt. Vernon 1 were evaluated at the early jointing stage, while others were evaluated at the flowering stage.

### Leaf rust response

Green Hammer was highly resistant to *Pt* races KFBJG, MCTNB, MGPSB, MHDSB, MJBJG, TCGJG, TCRDG, and TFTSB, moderately resistant to MNPSD and MPPSD, and susceptible to TBBGS and TNBJS (Table 2; Supplementary Table S5). Lonerider was susceptible to most *Pt* races, except TFTSB (Table 2). Gene postulation in Green Hammer based on the reactions of 20 Thatcher near-isogenic lines to the tested *Pt* races, suggested the presence of *Lr21* and *Lr39*. Green Hammer showed high resistance to *Pt* races that were avirulent to *Lr21* and *Lr39* (Table 2). Frequency distributions of linearized IT at the seedling stage ranged from 0 to 9 in the DH population with the presence of transgressive segregants with relatively high resistance to race MNPSD compared to the resistant parent Green Hammer (Fig. 2; Supplementary Table S6). Significant positive correlations were observed among seedling IT reactions to the seven *Pt* races, except for MNPSD (Supplementary Fig. S2).

**Fig. 2.**
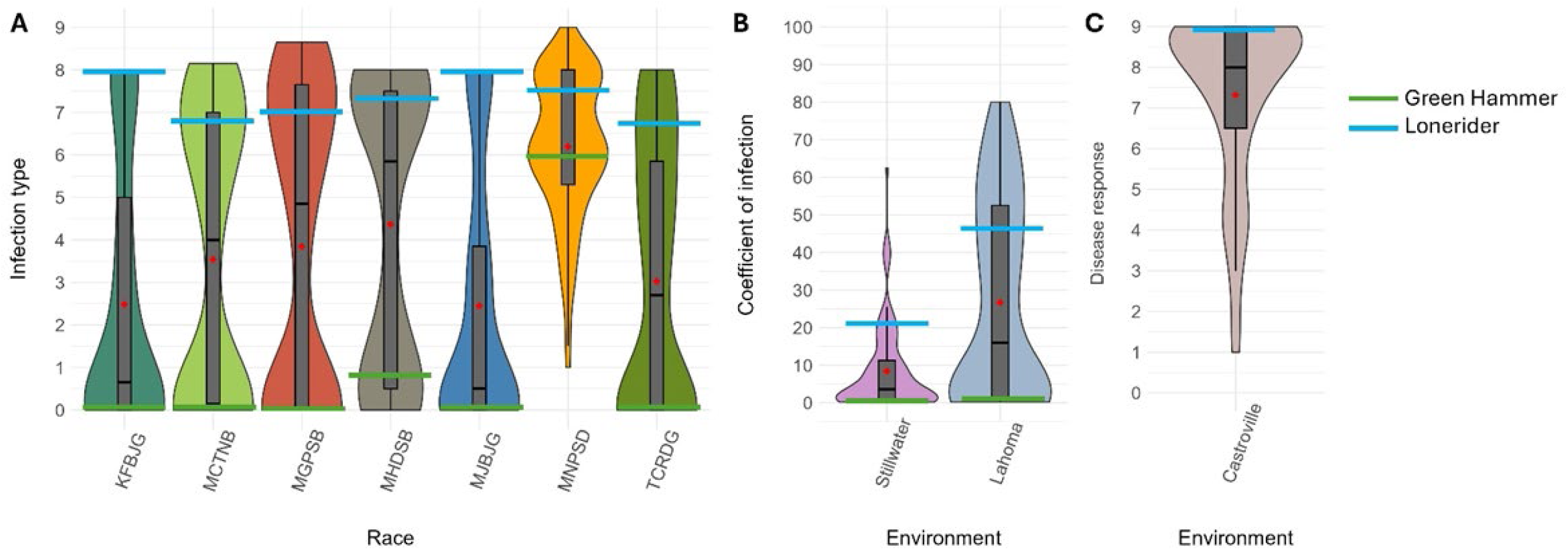
Distribution of leaf rust responses of 109 doubled haploid lines from the Green Hammer × Lonerider cross, including **A**, seedling infection types against seven *Puccinia triticina* races, **B**, coefficient of infection, and **C**, disease response at the adult plant stage across field environments. In the boxplots, black bold horizontal lines represent the medians and red diamonds represent the means. Green and blue horizontal lines indicate the mean leaf rust responses of the parents, Green Hammer and Lonerider, respectively.

**TABLE 2.**
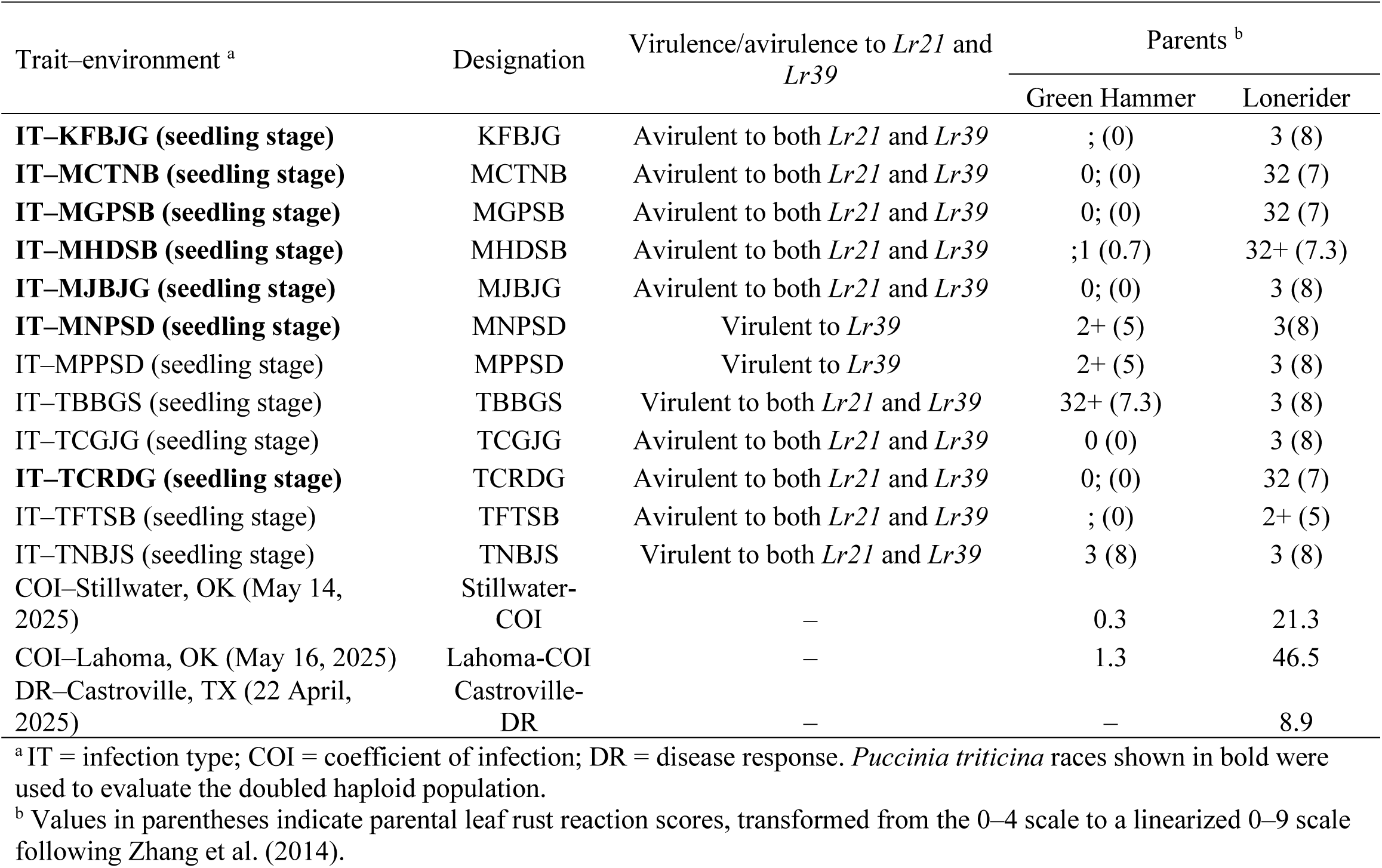
Leaf rust responses of Green Hammer and Lonerider at the seedling stage against 12 *Puccinia triticina* races and at the adult plant stage in field environments.

At the adult plant stage in field nurseries, the DH population showed a mean COI that ranged from 8.5 in Stillwater, OK to 26.7 in Lahoma, OK, with a lower disease pressure in Stillwater (Fig. 2; Supplementary Table S7). In Castroville, TX, the mean DR on a scale of 0-9 was 7.3, and the distribution was skewed toward susceptibility (Fig. 2). Leaf rust responses were positively and significantly correlated among field environments and between field environments and seedling responses to the seven *Pt* races (Supplementary Fig. S2).

### Genetic linkage map

After removing redundant markers and markers that deviated from the expected 1:1 segregation ratio at *p* ≤ 0.01, a total of 590 polymorphic markers were retained for QTL mapping, including 241 on the A genome, 235 on the B genome, and 114 on the D genome. No DH lines were removed for excessive crossover counts, and no markers were discarded for switched alleles or double crossovers. The final linkage map spanned 2,991.6 cM across 32 linkage groups representing all 21 wheat chromosomes (Supplementary Tables S8 and S9). However, no linkage groups were obtained for chromosome arms 1DL and 3BS. Chromosomes 2B, 2D, 3D, 4B, 7A, 7B, and 7D each formed two linkage groups, whereas chromosomes 1B and 5D each formed three linkage groups. All remaining chromosomes were represented by a single linkage group. Overall marker density was one marker per 5.07 cM. Individual linkage group length ranged from 3.2 cM for linkage group 7D-1, which contained only three markers, to 219.6 cM for linkage group 5A, which contained 41 markers.

### QTL analysis of stripe rust resistance

The QTL analysis identified 14 loci for stripe rust resistance in the DH population. Of these, 4 were major QTL, namely *QYr.osughln-2AS*, *QYr.osughln-2AL.1*, *QYr.osughln-2AL.2*, and *QYr.osughln-2BS.2*, whereas the remaining 10 had minor effects (Table 3). Among these four major QTL, *QYr.osughln-2AS* was the only major QTL contributed by Green Hammer and was detected in all locations except Rossville (Table 3; Fig. 3). It mapped to the short arm of chromosome 2A at 18.4–37.6 Mb based on IWGSC RefSeq v2.1 and explained 15.1%–94.7% of the phenotypic variation. The LOD scores for *QYr.osughln-2AS* ranged from 5.1 for 2025 Pullman DS to 69.6 for 2024 Mt. Vernon 1 IT. The SNP marker *S2A_18460882* was the closest marker to this QTL.

**Fig. 3.**
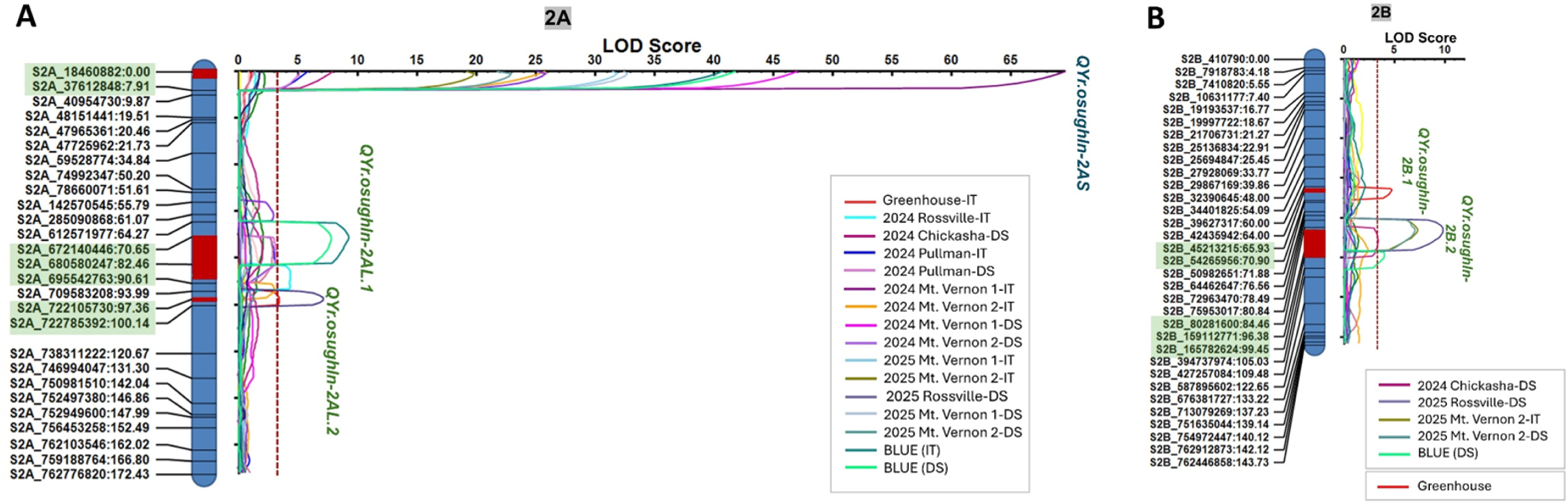
Genetic maps and inclusive composite interval mapping results for two chromosomes, **A**, chromosome 2A and **B**, chromosome 2B, carrying major quantitative trait loci (QTL) for adult plant stripe rust resistance in the Green Hammer × Lonerider population. The genetic map is shown in centiMorgans, and markers highlighted in light green represent the flanking markers for each identified QTL. Red bars on the genetic map indicate QTL regions. The red dashed line on the logarithm of odds (LOD) curves indicates the LOD threshold for QTL detection, which was set at 3.2 based on 1,000 permutations with p = 0.05. QTL shown in blue originated from Green Hammer, whereas QTL shown in green originated from Lonerider. IT = infection type; DS = disease severity (%); BLUE = multi-environment best linear unbiased estimates

**TABLE 3.**
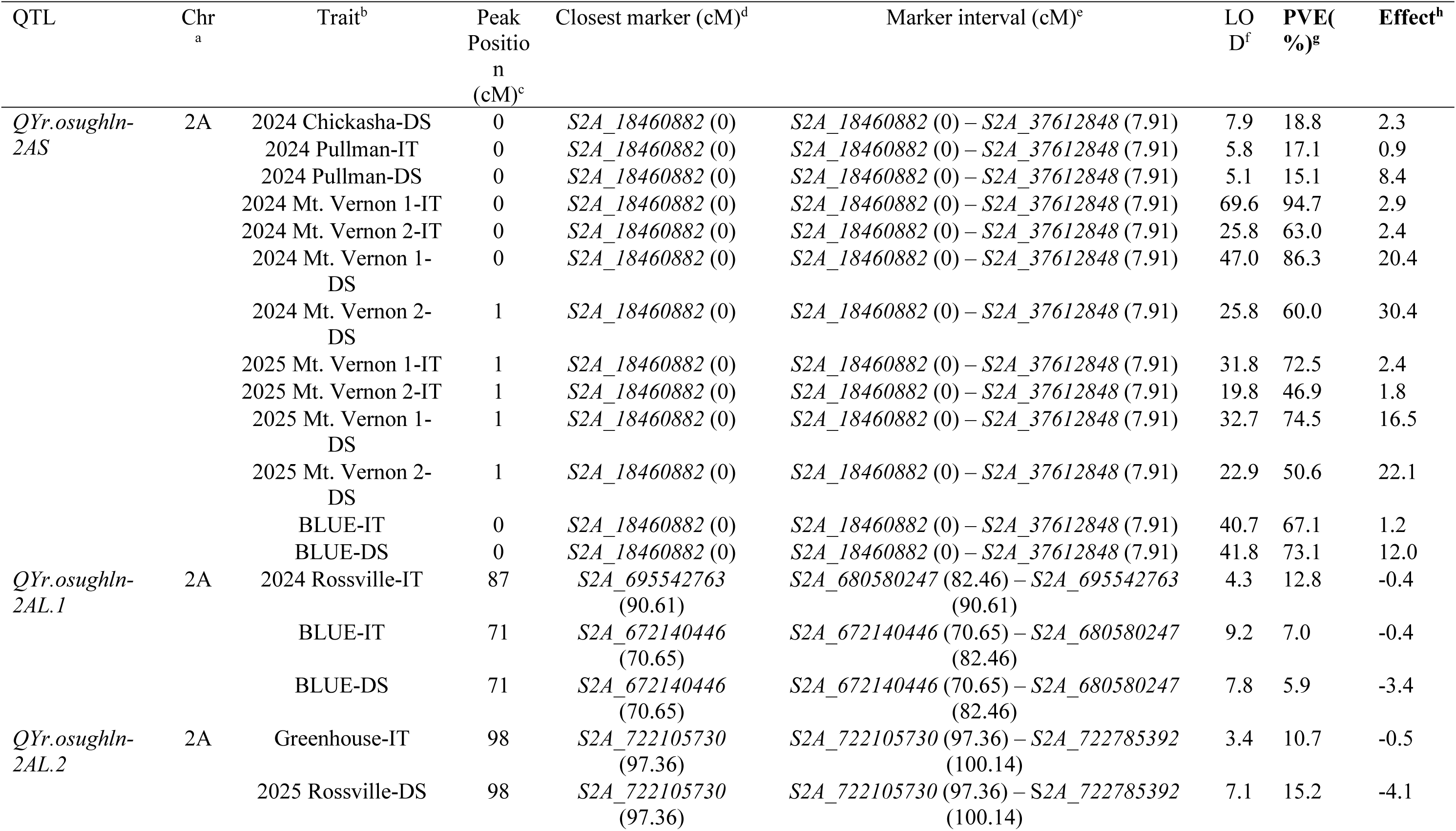
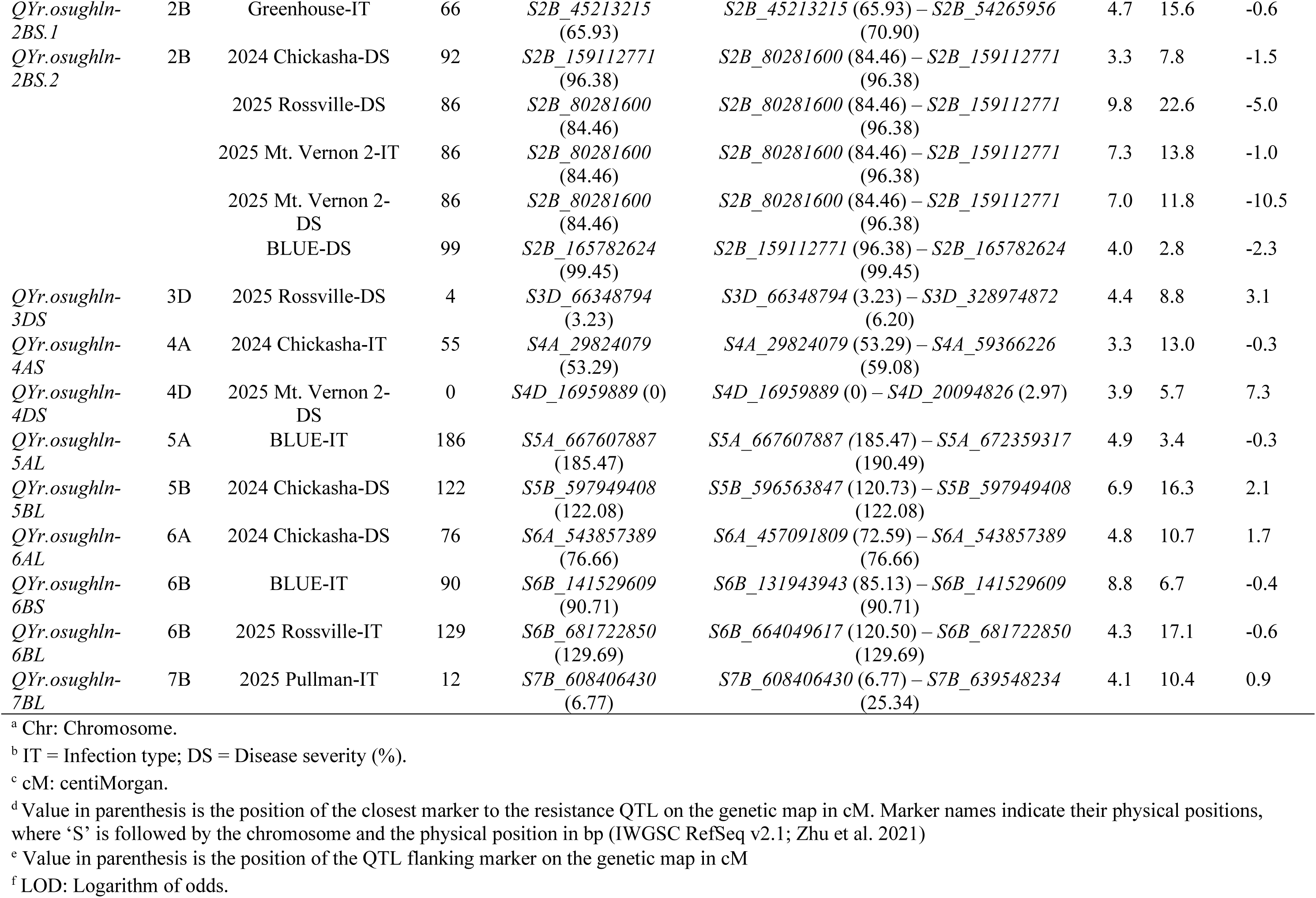
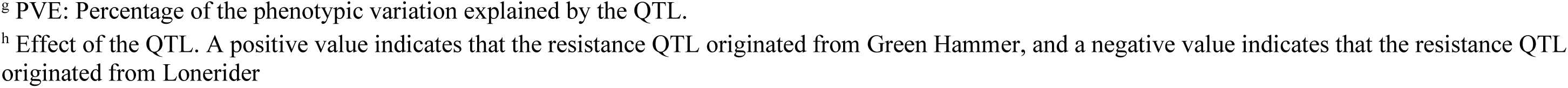
Summary of adult plant stripe rust resistance QTL detected in the Green Hammer × Lonerider doubled haploid population across environments.

Three major QTL were contributed by Lonerider, including *QYr.osughln-2AL.1* mapped to chromosome 2AL at 672.1–695.5 Mb and explained up to 12.8% of the phenotypic variation. It was detected with the 2024 Rossville IT and BLUE data (Table 3; Fig. 3). The second major QTL from Lonerider, *QYr.osughln-2AL.2*, was also mapped on the long arm of chromosome 2A at 722.1–722.7 Mb. It explained 10.7%–15.2% of the phenotypic variation, with LOD scores ranging from 3.4 to 7.1. The third major QTL from Lonerider, *QYr.osughln-2BS.2*, was identified on the short arm of chromosome 2B at 80.2–165.7 Mb. It was detected with the 2024 Chickasha DS, 2025 Rossville DS, 2025 Mt. Vernon 2 IT, 2025 Mt. Vernon 2 DS, and BLUE DS. This QTL explained up to 22.6% of the phenotypic variation, and its LOD scores ranged from 3.3 to 9.8 (Table 3; Fig. 3).

In total, 10 minor QTL were identified, with 5 contributed by Green Hammer and 5 by Lonerider (Table 3). The minor QTL from Green Hammer included *QYr.osughln-3DS*, *QYr.osughln-4DS*, *QYr.osughln-5BL*, *QYr.osughln-6AL*, and *QYr.osughln-7BL*, which were detected with the 2025 Rossville DS, 2025 Mt. Vernon 2 DS, 2024 Chickasha DS, 2024 Chickasha DS, and 2025 Pullman IT, respectively. The minor QTL contributed by Lonerider included *QYr.osughln-2BS.1*, *QYr.osughln-4AS*, *QYr.osughln-5AL*, *QYr.osughln-6BS*, and *QYr.osughln-6BL*, which were detected with the greenhouse IT, 2024 Chickasha IT, BLUE IT, BLUE IT, and 2025 Rossville IT data, respectively (Table 3).

### QTL analysis for leaf rust resistance

Eight QTL were identified for leaf rust responses, including 3 QTL contributed by Green Hammer and 5 by Lonerider (Table 4). Green Hammer contributed 2 major QTL, *QLr.osughln-1DS* and *QLr.osughln-2DS.1*, and 1 minor QTL, *QLr.osughln-2AS*. Lonerider contributed 1 major QTL, *QLr.osughln-2DS.2*, and 4 minor QTL, *QLr.osughln-2BS*, *QLr.osughln-3BL*, *QLr.osughln-3DS*, and *QLr.osughln-6DL*. Among the Green Hammer QTL, *QLr.osughln-1DS* was a major QTL mapped to the short arm of chromosome 1DS at 0.4–2.9 Mb and explained 2.0%–59.5% of the phenotypic variation. This ASR QTL was detected for seedling responses to *Pt* races KFBJG, MGPSB, MJBJG, MNPSD, and TCRDG and for adult plant responses in field environments Stillwater COI, Lahoma COI, and Castroville DR (Table 4; Fig. 4). The LOD scores for *QLr.osughln-1DS* ranged from 3.3 for MGPSB to 30.4 for Lahoma COI. This QTL was flanked by markers *S1D_465878* (0 cM) and *S1D_2922807* (6.13 cM), with the former being the closest marker. The second major QTL from Green Hammer, *QLr.osughln-2DS.1*, was mapped on chromosome 2DS at 1.6–15.6 Mb with peak marker *S2D_1647723* (Table 4; Fig. 4). This ASR QTL was detected for *Pt* races KFBJG, MCTNB, MGPSB, MHDSB, MJBJG, and TCRDG and for the Lahoma COI field environment. It explained up to 85.0% of the phenotypic variation, with LOD scores ranging from 5.8 to 50.9. The minor QTL from Green Hammer, *QLr.osughln-2AS*, was mapped to the short arm of chromosome 2A at 18.4–37.6 Mb and explained up to 6.4% of the phenotypic variation (Table 4). This QTL was detected at the seedling stage for *Pt* races KFBJG and MJBJG.

**Fig. 4.**
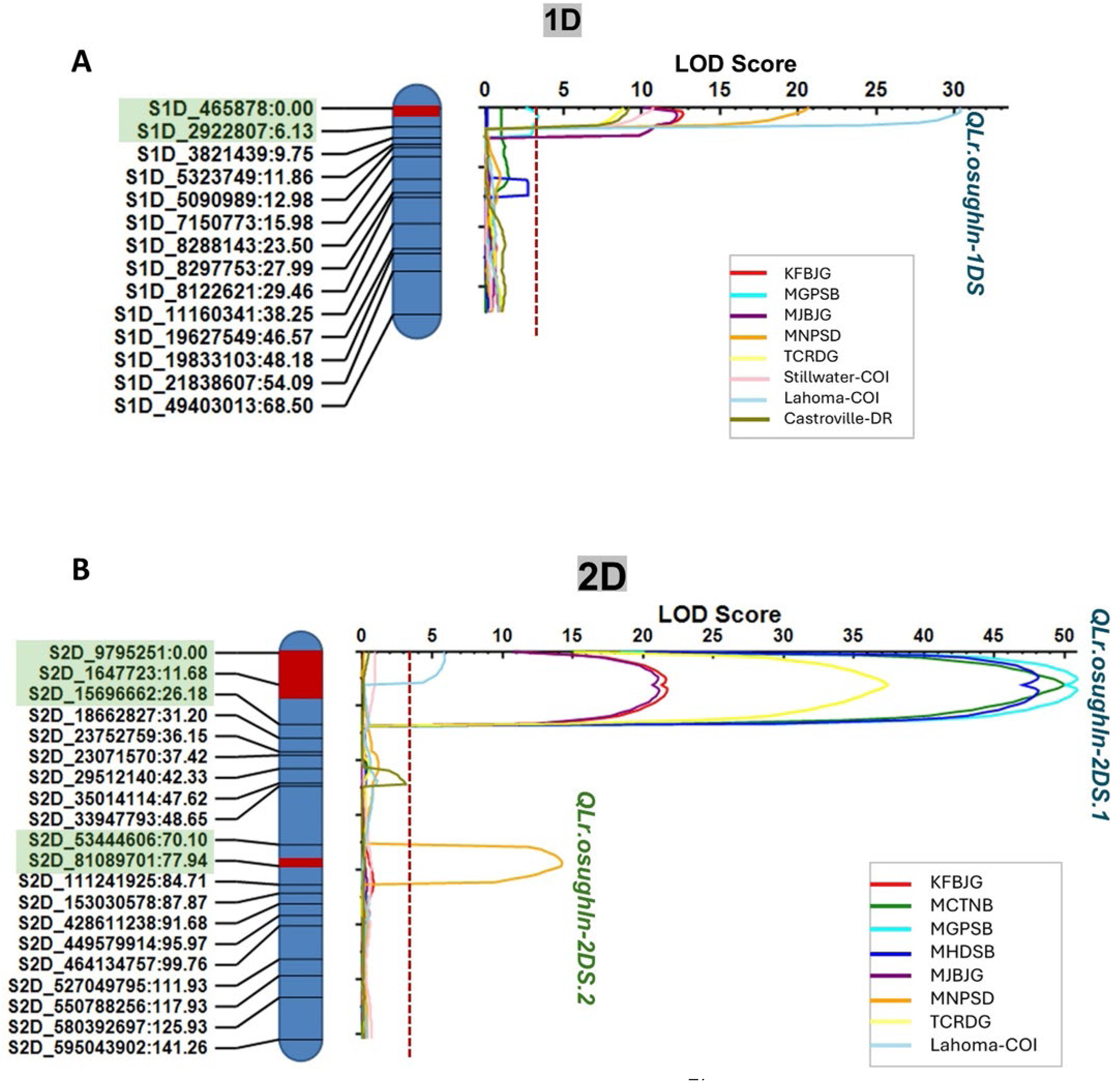
Genetic maps and inclusive composite interval mapping results for two chromosomes, **A**, chromosome 1D and **B**, chromosome 2D, carrying major quantitative trait loci (QTL) for leaf rust resistance in the Green Hammer × Lonerider population. The genetic map is shown in centiMorgans, and markers highlighted in light green represent the flanking markers for each identified QTL. Red bars on the genetic map indicate QTL regions. The red dashed line on the logarithm of odds (LOD) curves indicates the LOD threshold for QTL detection, which was set at 3.2 based on 1,000 permutations with p = 0.05. QTL shown in blue originated from Green Hammer, whereas QTL shown in green originated from Lonerider. COI = coefficient of infection; DR = disease response.

**TABLE 4.**
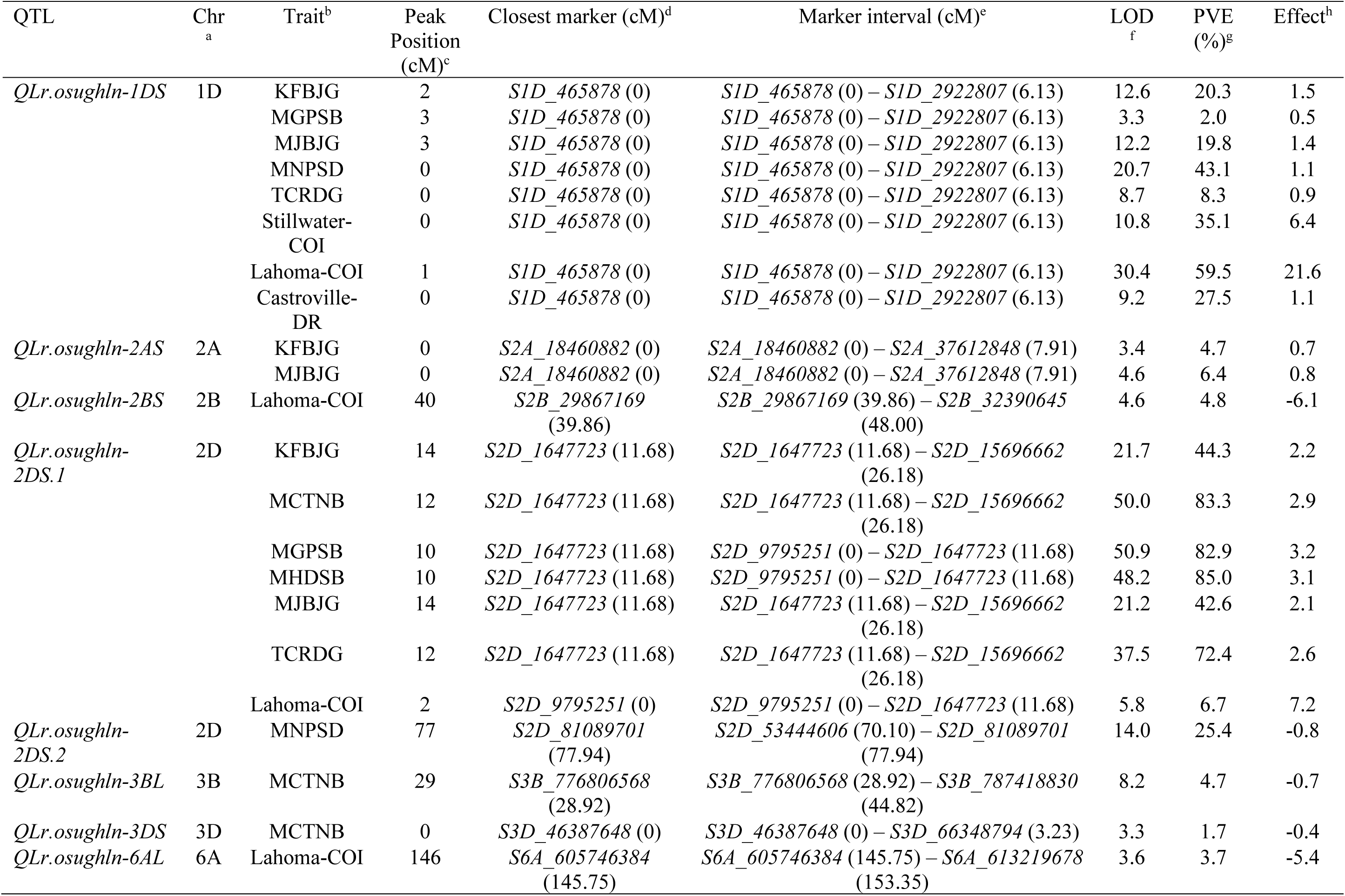
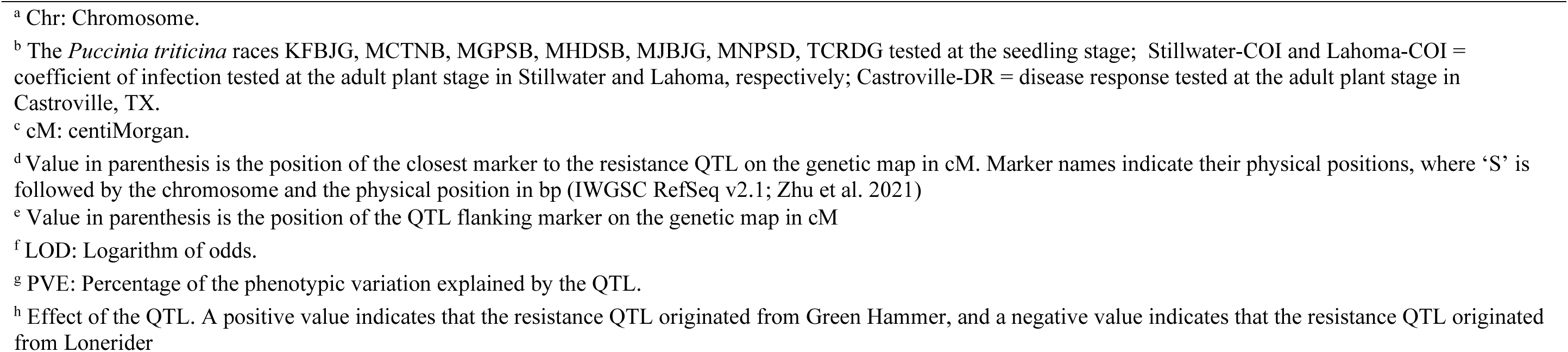
Summary of leaf rust resistance QTL detected in the doubled haploid population Green Hammer × Lonerider.

For Lonerider, the major QTL *QLr.osughln-2DS.2* was detected for seedling response to *Pt* race MNPSD (Table 4; Fig. 4). This QTL mapped to chromosome 2DS at 53.4–81.0 Mb and explained 25.4% of the phenotypic variation. The DH lines carrying *QLr.osughln-2DS.2* in combination with *QLr.osughln-1DS* showed significantly higher resistance to MNPSD than lines carrying *QLr.osughln-1DS* alone (Supplementary Fig. S3). Two minor ASR QTL, *QLr.osughln-3BL* and *QLr.osughln-3DS*, were detected for seedling response to MCTNB. They were mapped to chromosomes 3BL and 3DS and explained 4.7% and 1.7% of the phenotypic variation, respectively (Table 4). Two additional minor QTL from Lonerider were detected at the adult plant stage in Lahoma, namely *QLr.osughln-2BS* and *QLr.osughln-6DL*. These QTL mapped to chromosomes 2BS and 6DL, respectively, and explained up to 4.8% of the phenotypic variation (Table 4).

## Discussion

The virulence evolution of *Pst* races adapted to warmer climates and the high diversity of *Pt* races make stripe rust and leaf rust among the most economically important diseases of wheat globally (Beddow et al. 2015; Hovmøller et al. 2011; Kolmer and Fajolu 2022). Identification and characterization of rust resistance genes in modern cultivars allow their rapid deployment in breeding programs due to minimal linkage drag (Xu et al. 2025). In this study, we identified genes for stripe rust APR and leaf rust ASR in the bi-parental population Green Hammer × Lonerider. Green Hammer was evaluated in SRPN 2017, and based on the molecular markers used, it carries *Lr21*, *Lr37*/*Yr17*, *Lr77*, and *Lr78*. Leaf rust gene postulation also suggested the presence of *Lr21*, which is an ASR gene originating from the wild wheat relative *Aegilops tauschii* (Huang et al. 2003; Rowland and Kerber 1974). In the present study, the QTL *QLr.osughln-1DS* was found to be *Lr21*, which was effective against most tested *Pt* races, except TBBGS and TNBJS. This QTL was detected for *Pt* races avirulent to *Lr21* and across field environments. However, it was not detected with MCTNB and MHDSB, which are also avirulent to *Lr21*. This may be because another major ASR QTL *QLr.osughln-2DS.1* explained up to 85% of the phenotypic variation and masked *QLr.osughln-1DS*. *Lr21* was the first wheat rust resistance gene successfully cloned in 2003 and is located on chromosome 1DS at approximately 0.44 Mb (Huang et al. 2003). It is annotated as *TraesCS1D03G0002900.1*, which encodes a NLR protein (Huang et al. 2003; Tong et al. 2024). Gene postulation in the present study, along with molecular marker data from SRPN 2017, also supported the presence of *Lr21* in Green Hammer. Therefore, *QLr.osughln-1DS* in Green Hammer is likely *Lr21*. *Pt* races virulent to *Lr21* were first detected in 2010 (Kolmer and Anderson 2011). However, virulence to *Lr21* may be associated with reduced fitness and competitive ability in *P. triticina*, which may have contributed to the low frequency of *Lr21*-virulent races in the US and Canada, based on annual leaf rust surveys. Lakkakula et al. (2025) reported that *Lr21* was present at a low frequency, approximately 10%, among hard winter wheat (HWW) cultivars from the U.S. Great Plains based on molecular markers. Together, the low frequency of *Lr21* in contemporary cultivars and the potential fitness penalty in the pathogen associated with *Lr21* virulence suggest that *Lr21*, and Green Hammer carrying *Lr21*, remain valuable sources of leaf rust resistance for future breeding programs.

The *Lr* postulation also suggested the presence of *Lr39* in Green Hammer. *Lr39* is an ASR gene transferred from *Ae. tauschii* accession TA1675 into common wheat and was mapped on chromosome 2DS near the SSR marker *gdm35* at approximately 13.9 Mb (Singh et al. 2004). Subsequently, *Lr41*, another ASR gene mapped to a similar region, was reported to be allelic or tightly linked to *Lr39*. In our study, QTL analysis identified a major QTL, *QLr.osughln-2DS.1*, mapped to 1.6–15.6 Mb, which overlaps the reported genomic region of *Lr39*. This QTL was highly effective against *Pt* races avirulent to *Lr39* at the seedling stage and was also detected in the Lahoma field. Other *Lr* genes have also been reported in the genomic region of *QLr.osughln-2DS.1*, including *Lr22a* and *Lr80*. *Lr22a* is an APR gene introgressed from *Ae. tauschii* and linked to the SSR marker *gwm296* (16.7 Mb) (Hiebert et al. 2007). *Lr80* is an ASR gene mapped in the Indian landrace Hango-2 and flanked by KASP markers *KASP_17425* at 6.8 Mb and *KASP_17148* at 10.5 Mb (Kumar et al. 2021). However, based on gene postulation and QTL position, *QLr.osughln-2DS.1* is most likely *Lr39*. Virulent races to *Lr39* occur at a relatively high frequency in the US Great Plains (Kolmer and Fajolu 2022). Despite this, the interaction between *QLr.osughln-1DS*, which represents *Lr21*, and *QLr.osughln-2DS.1*, which represents *Lr39*, was significant for leaf rust responses to most *Pt* races and in the Stillwater and Lahoma fields. Among the seven *Pt* races tested on the DH population, MNPSD was the only race that was virulent to *Lr39* and avirulent to *Lr21*. Green Hammer showed a moderately resistant reaction to MNPSD (IT = 2+), whereas it showed high resistance to the other tested races avirulent to *Lr21* and *Lr39* (IT = ; to ;1). This pattern suggested that the combined effects of *Lr21* and *Lr39* provide higher levels of resistance than *Lr21* alone. In addition, a major-effect QTL, *QLr.osughln-2DS.2*, contributed by Lonerider, was detected for responses to MNPSD. Although Lonerider was susceptible to MNPSD, DH lines carrying only this QTL showed improved resistance compared with Lonerider, indicating a positive effect of this locus outside the Lonerider genetic background. Furthermore, this QTL likely explains the presence of transgressive segregants with higher resistance to MNPSD than that of Green Hammer.

Lonerider was evaluated in SRPN in 2016 and 2017 and was reported to carry *Lr77*. Similarly, Green Hammer was also reported to carry *Lr77* based on molecular markers. *Lr77* is an APR gene originally identified in the wheat cultivar ‘Santa Fe’ on chromosome 3BL and is closely linked to the SNP marker *IWB10344* at 769.5 Mb (Kolmer et al. 2018b). *Lr77* was later identified in the OSU HRWW cultivar ‘Duster’ (Kolmer et al. 2019) and is widely present in contemporary Great Plains HRWW (Kolmer et al. 2018b; Lakkakula et al. 2025). We did not detect *Lr77* because this locus was fixed in this population, as it is present in both parents. In addition, marker data suggested that Green Hammer carries *Lr78*. *Lr78* is an APR gene derived from the Brazilian wheat cultivar ‘Toropi’ and was mapped on chromosome 5DS and linked to the SNP marker *IWA6289* at 33.0 Mb (Kolmer et al. 2018a). In this study, no QTL was detected in the genomic region of *Lr78*. This may reflect limited diagnostic power of the *Lr78*-linked marker (Kolmer et al. 2018a), leading to a false-positive call in Green Hammer, or masked *Lr78* partial effect by the major gene *Lr21*.

The major stripe rust APR QTL, *QYr.osughln-2AS* identified in Green Hammer, was consistently associated with stripe rust responses across all environments except in the greenhouse and Rossville, KS. *QYr.osughln-2AS* was physically located on chromosome 2AS at 18.4–37.6 Mb, within the genomic region of the 2N^v^S translocation from *Ae. ventricosa*. The 2N^v^S segment, which is approximately 33 Mb, was originally introduced into the French wheat cultivar ‘VPM1’ for eyespot resistance and was later found to carry the *Lr37*/*Yr17*/*Sr38* rust resistance gene cluster (Gao et al. 2021). In the present study, SNP markers spanning 0.50 Mb to 33.2 Mb co-segregated. Among these markers, *S2A_18460882* was retained for linkage mapping, whereas the remaining co-segregating markers were excluded. Marker *S2A_18460882* was closely linked to *QYr.osughln-2AS*. The Kompetitive allele-specific PCR (KASP) marker (*Lr37/Yr17/Sr38_GBG-KASP*) on the 2N^v^S (Li et al. 2023b) indicated the presence of this translocation in Green Hammer, which carries *Lr37*/*Yr17/Sr38*. Together, these results support that *QYr.osughln-2AS* is associated with resistance on or linked to the 2N^v^S translocation. The source of the 2N^v^S introgression in Green Hammer is likely the HRWW cultivar ‘OK Bullet’. OK Bullet has ‘Jagger’ in its pedigree, and 2N^v^S was first introduced into the HWW Jagger and subsequently used extensively in wheat breeding programs for rust resistance (Gao et al. 2021). This translocation, has been reported at a high frequency in contemporary US HWW (Lakkakula et al. 2025; Mu et al. 2020; Sharma et al. 2025). In addition to the *Lr37*/*Yr17*/*Sr38* gene cluster, the 2N^v^S segment has been shown to carry other rust resistance loci, including a high-temperature adult plant (HTAP) resistance gene, *YrM1225* (Li et al. 2023b). *QYr.osughln-2AS* was identified in almost all field environments, but its effect was not significant when the DH population was tested at cool temperatures in the greenhouse (4–20°C), suggesting that *QYr.osughln-2AS* is likely the HTAP resistance gene *YrM1225*. Although the major QTL *QYr.osughln-2AS* was not detected in the greenhouse at cool temperatures, Green Hammer remained resistant. This suggests that Green Hammer carries other undetected QTL that could be fixed in the DH population because of shared resistance QTL between Green Hammer and Lonerider.

A minor effect QTL, *QLr.osughln-2AS* identified from Green Hammer for seedling leaf rust responses to *Pt* races KFBJG and MJBJG, was flanked by the same markers and shared the same peak marker as *QYr.osughln-2AS*. The ASR gene *Lr17a* is tightly linked to *Lr37*, and *Pt* virulence profiles are often similar for *Lr17a* and *Lr37* (Bremenkamp-Barrett et al. 2008; Xue et al. 2018). Most contemporary *Pst* and *Pt* races are virulent to *Yr17* and *Lr17a/Lr37*, respectively (Kolmer and Fajolu 2022; Wang et al. 2022). *QLr.osughln-2AS* was detected only for races against which *Lr17a/Lr37* is effective, except for TCRDG. The lack of detection for TCRDG likely reflects masking by two other effective QTL, *QLr.osughln-1DS* (*Lr21*) and *QLr.osughln-2DS.1* (*Lr39*). Therefore, *QLr.osughln-2AS* is most likely *Lr17a/Lr37*.

Three major QTL were identified in Lonerider for stripe rust APR. *QYr.osughln-2AL.1* (672.1–695.5 Mb), which was detected for 2024 Rossville IT and BLUE, and *QYr.osughln-2AL.2* (722.1–722.7 Mb), which was detected for the greenhouse assay and 2025 Rossville DS, were located adjacent to each other on chromosome 2AL. The genomic interval for *QYr.osughln-2AL.2* would extend to 711.9 Mb if redundant markers were included, resulting in an effective interval of 711.9–722.7 Mb. Several previously reported APR genes or QTL on chromosome 2AL were within the genomic region of QTL identified in Lonerider, including *Yr86* (710.3–712.6 Mb) from the Chinese wheat cultivar Zhongmai 895 (Cao et al. 2024; Zhu et al. 2023), *Yrxy2* linked to *Xbarc5* at 678.1 Mb (Zhou et al. 2011), *QYrWS.wgp-2AL* (611.6–684.7 Mb) (Upadhaya et al. 2025), and *QYrPI197734.wgp-2A* (559.7–713.4 Mb) (Liu et al. 2020). Another major effect QTL contributed by Lonerider, *QYr.osughln-2BS.2*, was detected for stripe rust responses in Chickasha, Rossville, and Mt. Vernon 2. It mapped to chromosome 2BS at 80.2–165.7 Mb. Several APR QTL have also been reported within the genomic region of *QYr.osughln-2BS.2*, including *QYr.sgi-2B.1* linked to *gwm148* at 108.4 Mb (Ramburan et al. 2004), *QYr.hebau-2BS* at 134.5 Mb (Gebrewahid et al. 2020), and *Qyrlov.nwafu-2BS* flanked by *IWA5377* (147.6 Mb) and *IWA5830* (173.0 Mb) (Wu et al. 2017).

This study identified and characterized the genetic basis of APR to stripe rust and ASR to leaf rust in the bi-parental population Green Hammer × Lonerider. The major QTL, *QYr.osughln-2AS*, mapped to the 2N^v^S translocation accounted for most of the stripe rust APR in Green Hammer. For leaf rust, ASR was mainly conferred by the major QTL, *QLr.osughln-1DS* and *QLr.osughln-2DS.1*, mapped near *Lr21* and *Lr39*, respectively. Lonerider also carried major effect APR QTL to stripe rust on chromosomes 2AL and 2BS. In this study, we identified previously designated genes as well as unknown genes that can be used in breeding programs to enhance leaf rust and stripe rust resistance in wheat breeding programs.

## Acknowledgements

This project was funded by the US Department of Agriculture National Institute of Food and Agriculture Grant # 2024-67013-42587. The mention of trade names or commercial products in this publication is solely to provide specific information and does not imply recommendation or endorsement by the US Department of Agriculture. The USDA is an equal opportunity provider and employer.

## Data Availability Statement

All data generated or analyzed during this study are included in this published article and its supplementary files submitted with this manuscript. The raw sequencing data can be accessed from the NCBI Short-Read Archive BioProject PRJNA1462017 (https://www.ncbi.nlm.nih.gov/sra/PRJNA1462017).

## Conflict of interest statement

The authors declare no conflicts of interest.

## Notes

### Competing Interest Statement

The authors have declared no competing interest.

