## Supplemental Figures for "Mapping of Stripe Rust and Leaf Rust Resistance Genes in the Hard Red Winter Wheat Population Green Hammer × Lonerider"

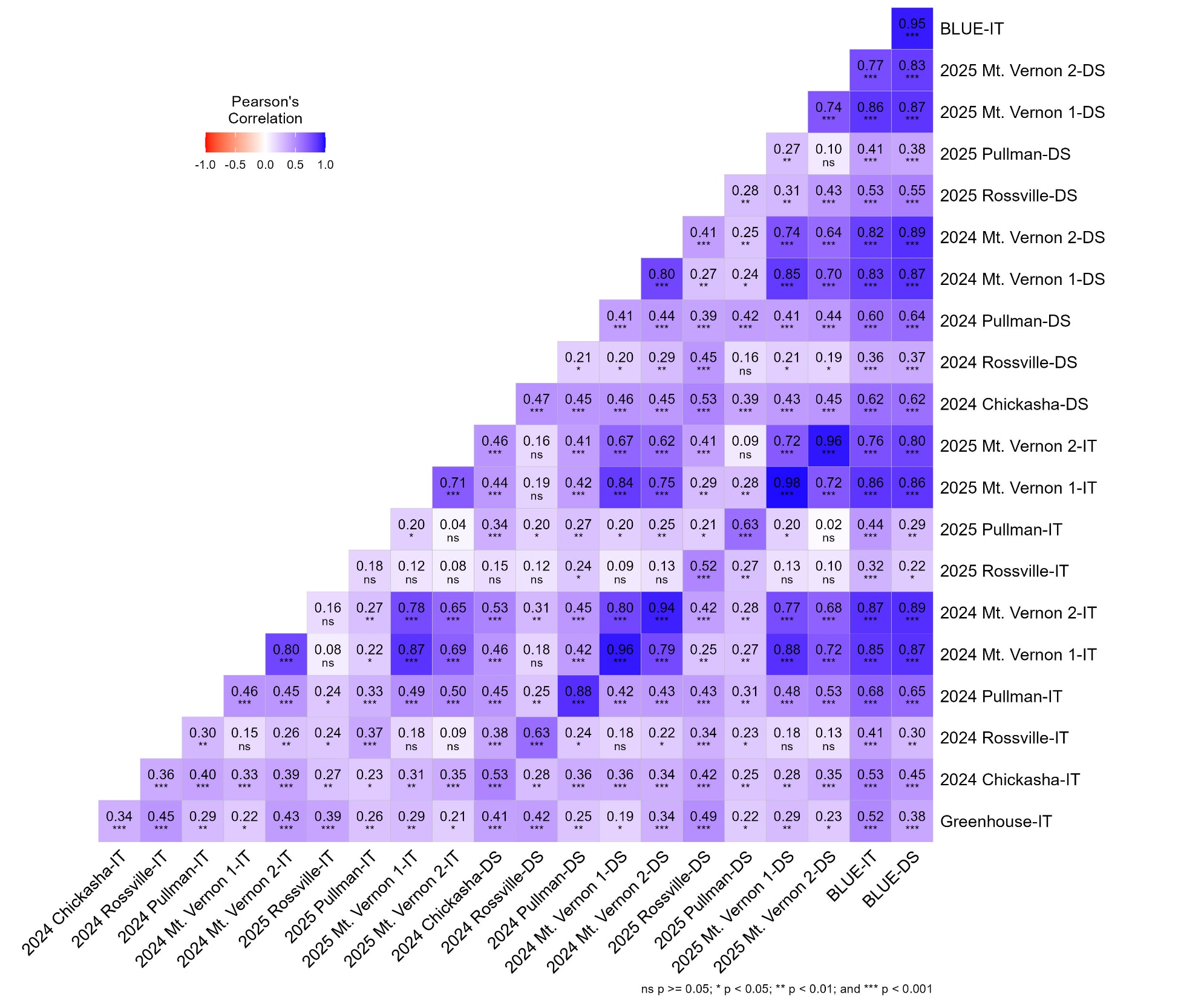

**Supplementary Fig. S1.** Correlations between stripe rust responses in different environments for the doubled haploid population Green Hammer × Lonerider. IT = infection type; DS = disease severity (%); BLUE = multi-environment best linear unbiased estimates.

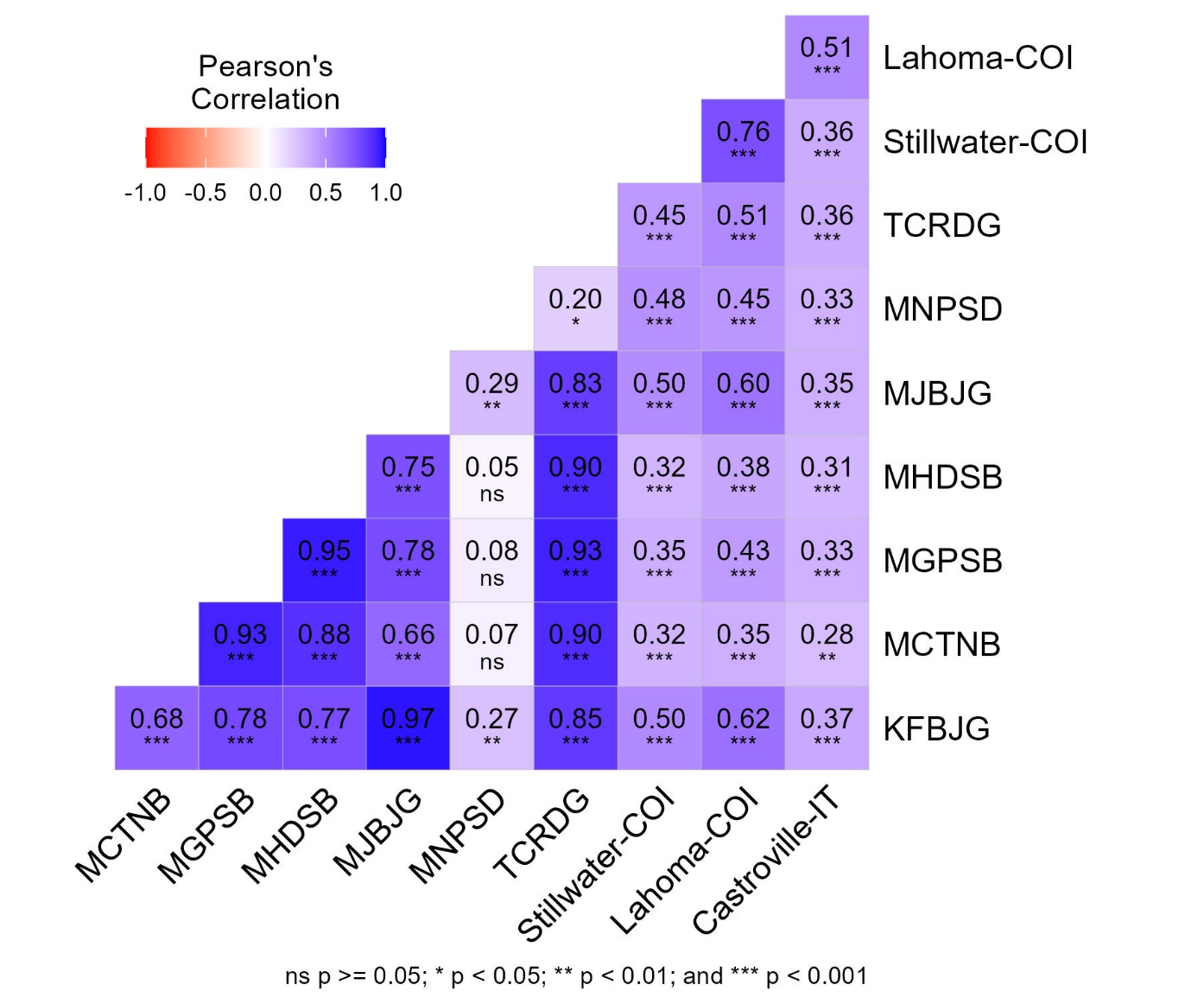

**Supplementary Fig. S2.** Correlations between leaf rust responses of 109 Green Hammer × Lonerider doubled haploid lines at the seedling stage against seven *Puccinia triticina* races and at the adult plant stage across different field environments. COI = coefficient of infection; IT = infection type.

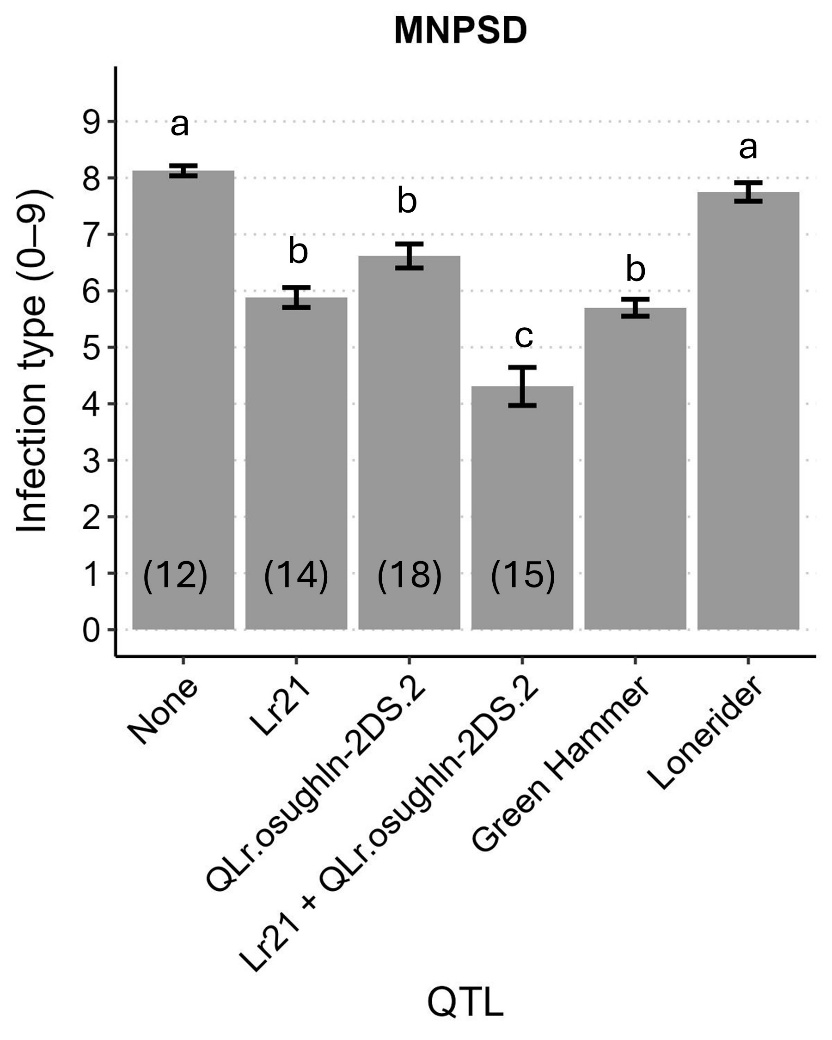

**Supplementary Fig. S3.** Frequency distribution of leaf rust responses to *Puccinia triticina* race MNPSD among doubled haploid lines carrying combinations of *QLr.osughln-1DS* (*Lr21*) from Green Hammer and *QLr.osughln-2DS.2* from Lonerider in the Green Hammer × Lonerider biparental population. Numbers beneath each bar indicate the number of lines carrying each QTL combination. Bars labeled with different letters differ significantly based on Tukey’s HSD test**.**
